# Global mosquito virome profiling and mosquito spatial diffusion pathways revealed by marker-viruses

**DOI:** 10.1101/2022.09.24.509300

**Authors:** Lu Zhao, Ping Yu, Chenyan Shi, Lijia Jia, Atoni Evans, Xiaoyu Wang, Qun Wu, Guodian Xiong, Zhaoyan Ming, Ferdinand Salazar, Bernard Agwanda, Dennis Bente, Fei Wang, Di Liu, Zhiming Yuan, Han Xia

## Abstract

Mosquitoes are vectors of numerous emergence and reemergence of mosquito-borne diseases, leading to an overwhelming global challenge. The boom in metagenomic studies promoted the increase of mosquito viruses being described, and studies from different regions of the world showed that mosquitoes harbor abundant and diverse viromes. However, there is still a lack of large-scale systematic comparison of viromes among various ecological factors such as the mosquito species/genera, location, etc, and consistent patterns associated with these factors are not clear. Here, we provide an overview of virome profiling that integrated the perspective of mosquito genus, locations, and climates based on the global scale, and redefined the ‘core-virome’. Our results also further implicate some mosquito-associated viruses strongly associated with the ecological factors and highlighted the evidence of mosquitoes’ cross-regional movements by the candidate marker viruses. The study may be helpful in gaining new insights into strategies to prevent arbovirus epidemics.

## 1. Introduction

Mosquitoes not only transmit arboviruses but are also naturally infected with a wide range of viruses, which are defined as insect-specific viruses (ISVs). In contrast to arboviruses, which cause human and animal emerging and reemerging infectious diseases and threaten regional and global public health, ISVs have a restricted host range, which can only infect and replicate in invertebrate cells, but is incapable to replicate in vertebrate cells or infecting humans and other vertebrates. As climate change intensifies, the acceleration of globalization and urbanization, and the expansion of human activities, geographic range of mosquito vectors are expending, which facilitated the spread of arboviruses[1]. Surveillance of mosquito vectors and the viruses they carried is an important tool to help raise the preparedness and prevention for mosquito-borne disease outbreaks. In the last decades, a growing number of metaviromic studies highlighted the abundance and diversity of viruses in mosquitoes[2–5]. The recent advances in viral metagenomics also facilitated the rapid growth of ISVs discovery and identified in mosquitoes[6].

Although ISV is rarely reported that it can cause disease and death of mosquitoes, the existence of this kind of virus affects some physiological functions of mosquitoes. And importantly, the stable and persistent existence of ISV in mosquito population as well as their continuous mutation and evolution, make it possible to expand their host range to infect humans and vertebrate animals[7–9]. In addition, many studies have shown that ISVs (e.g. Culex Flavivirus[10–12], Palm Creek virus[13], Cell fusing agent virus[14], Phasi Charoen-like phasivirus[14],) can interact with arboviruses in mosquito, promote or inhibit the replication and dissemination of arboviruses, and finally affect the transmissibility of arboviruses from mosquito to human or animal. Therefore, understanding the abundance and diversity of mosquito associated viruses (both arbovirus and ISVs) and their influencing factors on mosquitoes under different location, climate and environmental conditions, and determining their viral structure and core virus are of great significance for in-depth understanding of the role and function of these viruses.

There are a lot of research focused on the metavirome in mosquitoes by country or region, and these studies have expanded our understanding of the breadth of viral diversity present in mosquito populations, but limited knowledge is known about their genetic structure and ecological dynamics change by host species, through time and space, or by environmental factors through global perspective. Virome studies of mosquitoes collected from USA[15], Guadeloupe[16], Europe[17], China[18], and Australia[19] have revealed that mosquito species was the main factor that shapes the composition and diversity of virome, and stable species-specific ‘core-virome’ in mosquito populations were defined. The examination of the mosquito virome across different life stages of both lab-reared and wild-caught *Aedes albopictus* mosquitoes demonstrated the stability and vertically transmitted of the core-virome[20]. A recent longitudinal study investigated the mosquito viromes dynamic through a year in a certain location in Yunnan province, China[21]. Strong variance in viral compositions and abundance were observed for different seasons and host species, but not for gender. However, these studies are restricted to between two or just a few mosquito species within a particular country or district [22]. The meta-virome survey to combine all globally published mosquito virome data can draw a comprehensive map of the ISVs distribution, define more plausible core viruses and give a deep insight into the determined factors (e.g. host species, climate, location) for virome structure. Meantime, the SNPs of core viruses can let us simulate the migration of their mosquito host.

In this study, we analyzed 685 mosquito metagenomic sdatesets originating from 21 different countries across five continents. We provide an overview of virome profiling that integrated the perspective of mosquito genus, locations, and climates based on the global scale, and redefined the ‘core-virome’. Our results also further implicate that some mosquito-associated viruses strongly associated with the ecological factors and highlighted the evidence of mosquitoes’ cross-regional movements inferred by the geographical reconstruction of candidate marker viruses.

## 2. Results

### 2.1 Sampling information of global mosquito metagenomic datasets

To provide a broader overview of virome in mosquitoes, we generated 77 viral metagenomic data from three mosquito genera across six provinces in China, and three counties in Kenya acquired from 2014 to 2021. Moreover, 608 exploitable mosquito viral metagenomic datasets were selected and downloaded from the Gene Expression Omnibus (GEO) database (Supplementary File 1). These mosquito samples belonged to 8 mosquito genera included: *Culex* spp., *Aedes* spp., *Anopheles* spp., *Armigeres* spp., *Coquillettidia* spp., *Culiseta* spp., *Mansonia* spp., and *Psorophora* spp., and was predominantly *Culex* spp., *Aedes* spp., *Anopheles* spp. They were collected from 2008 to 2021 and were distributed across the five continents involving 21 different countries, and the climate between sampling regions was diverse and showed 16 different types (Figure 1).

**Figure 1.**
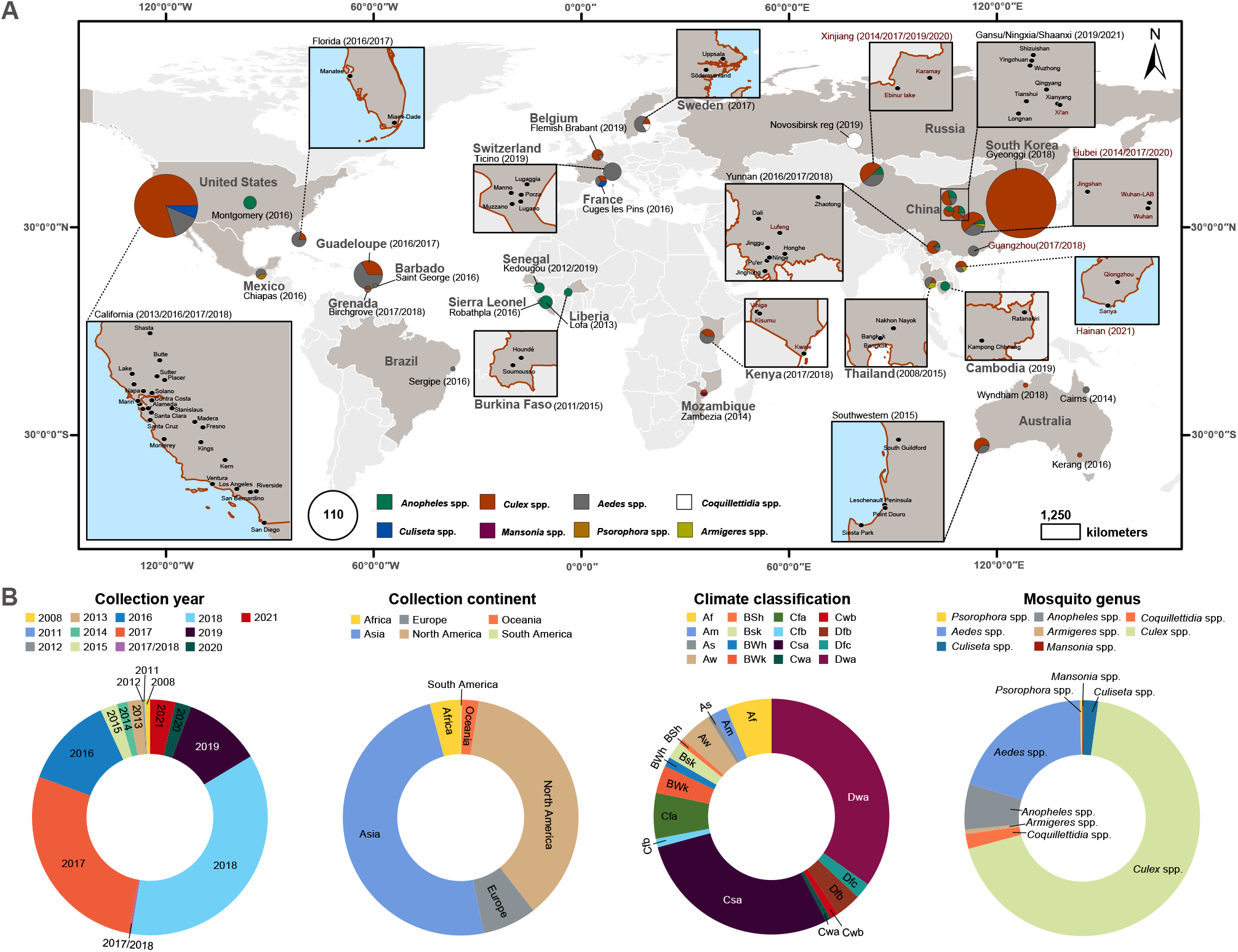
Sampling information of global viral metagenomic datasets. (A) Geographic distribution of mosquito samples. The labels in red sequencing data from our laboratory, and the label in black were sequencing data from public database. (B) The generalization of sample collection. From 2008 to 2021, mosquito samples involving 8 mosquito genera were collected in Africa, Asia, Europe, South America, and Oceania and distributed in 16 climate zones.

### 2.2 A peek of mosquito eukaryotic viromes

Here, we focused on the eukaryotic viruses in mosquitoes and filtered out bacteria, fungi, and protozoa-related viral nonredundant reads. And we mapped all clean reads back to the eukaryotic contigs, a viral species was considered present if its contig length > 500 bp and more than 500 reads mapped to it. According to the taxonomic annotation, 773 eukaryotic viral species were assigned across all samples. The eukaryotic viruses were identified into 58 viral families, 5 unclassified viral orders, and other unclassified viruses (including floating genera, such as Negevirus), The principal component analysis (PCA) based on different conditions showed that the differentiation of viridae among various conditions is higher than that of viral species among different conditions, which also suggested the similarity among viral families was better than that of viral species (Supplementary Figure 1, 2). The nucleic acid types of these single-stranded or segmented viral genomes include ssRNA (+/ -), ssRNA-RT, ssDNA (+/-), dsRNA and dsDNA, etc., Of these 58 viral families, there were only 13 viral species classified as mosquito-borne viruses, and they were mainly distributed in the families *Flaviviridae, Rhabdoviridae, Phenuiviridae, Peribunyaviridae, Reoviridae* and *Nodaviridae*. And *Flaviviridae, Togaviridae, Perbunyaviridae, Rhabdoviridae, Mesoniviridae, Tymoviridae, Birnaviridae, Nodaviridae, Reoviridae, Parvoviridae, Permutotetraviridae, Solemoviridae, Phasmaviridae, Iflaviridae, Orthomyxoviridae, Xinmoviridae* and *Totiviridae* were reported to contain members of mosquito-specific viruses. Negeviruses were also mosquito-specific viruses that have been isolated across the world. In addition to these known mosquito-associated viruses, the majority of them remain to be characterized and have not actually been isolated or identified (Figure 2). Although some studies have shown variations in virus diversity and abundance in different mosquitoes, there is still a lack of systematic comparison between mosquito species or genera. On a gross scale, *Aedes* spp. and *Anopheles* spp. have higher viral richness than *Culex* spp. whether the mean number of viral species or family, and *Armigeres* spp. showed the highest viral abundance, while *Psorophora* spp. has the lowest abundance (Supplementary Figure 3).

**Figure 2.**
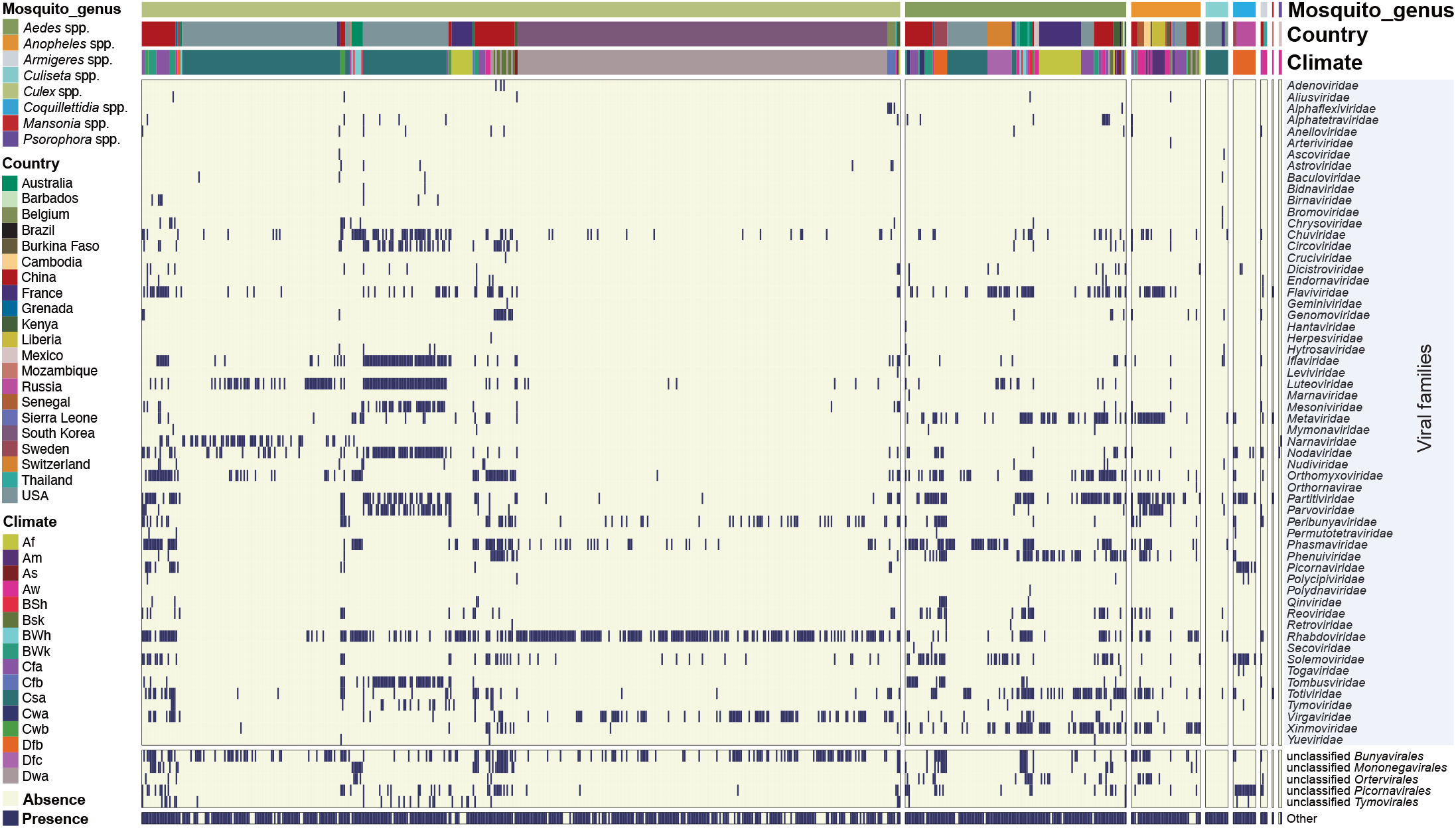
Classification of eukaryotic viruses in mosquitoes. The heatmap showing the presence of eukaryotic viruses. Eukaryotic viruses were considered present in the samples if its contig length was > 500 bp and > 500 reads mapped to it. All mosquito eukaryotic viruses were classified into 58 viral families, 5 unclassified viral orders, and other unclassified viruses. At the same time, each sample was annotated with its mosquito genera, location, and climate information.

To further compared abundance of the 58 eukaryotic viral families between the different mosquito genera. We counted the percentage of individual eukaryotic viral family reads / total eukaryotic viral reads in each sample, and the proportion of more than 20% in multiple (> 5) mosquito samples were regarded as the dominant family (Supplementary Figure 4). The results showed that the *Alphatetraviridae, Phenuiviridae, Metaviridae, Partitiviridae, Phasmaviridae, Solemoviridae* and *Totiviridae* were observed to be the dominant viral families of *Aedes* spp. samples. The *Culex* spp. samples contains 13 dominant viral families, which were *Iflaviridae, Luteoviridae, Mesoniviridae, Narnaviridae, Nodaviridae, Orthomyxoviridae, Partitiviridae, Parvoviridae, Phasmaviridae, Tombusviridae, Totiviridae, Virgaviridae* and *Rhabdoviridae*. There were 3 dominant viral families in *Anopheles* spp. samples including *Flaviviridae, Phasmaviridae* and *Xinmoviridae*. Unclassified viruses account for a relatively large proportion of the eukaryotic viruses in *Culiseta* spp. samples, *Armigeres* spp. samples as well. Among the *Coquillettidia* spp. samples, *Picornaviridae* was the dominant viral family, and viruses that classified as unclassified *Picornavirales* are highly abundant. The sample number of *Psorophora* spp. and *Mansonia* spp. were both less than 3, according to the existing data, *Nodaviridae* and *Narnaviridae* was more abundant in *Psorophora* spp. and most eukaryotic viruses in *Mansonia* spp. samples are unclassified viruses.

A further comparison and intersection among the viral species in each mosquito genus revealed the shared and specific eukaryotic viromes in mosquitoes. There were no shared viral species among the eight mosquito genera, but the most common mosquito genus included *Culex* spp., *Aedes* spp., and *Anopheles* spp. had enriched shared viral species (Figure 3A). Of these, the mosquito-associated viruses were distributed in 20 viral families, 4 unclassified viral orders, and 22 unclassified viruses, and most of them are insect-specific viruses (Figure 3B), which also contains arbovirus like Dengue virus, it suggested the potential for horizontal transmission of these viruses among different mosquitoes. Furthermore, except the *Mansonia* spp., the remaining 7 mosquito genus possess the mosquito-genera specific viruses. *Culex* spp. harbor the most numerous and abundant mosquito-genera specific viruses, which belong to 27 viral families, 2 unclassified viral orders, and 34 unclassified viruses. 47 mosquito-associated viruses were specifically observed in *Aedes* spp., One of them is Chikungunya virus,a mosquito-transmitted alphavirus belonging to the *Togaviridae* family. The Bluetongue virus was only seen in *Anopheles* spp., and the other mosquito-borne virus Sindbis virus was found to be associated only with *Coquillettidia* spp. The number in mosquito-genera specific virome of *Culiseta* spp., *Psorophora* spp., and *Armigeres* spp. was relatively smaller, and the non-mosquito-associated viruses accounted for a large proportion.

**Figure 3.**
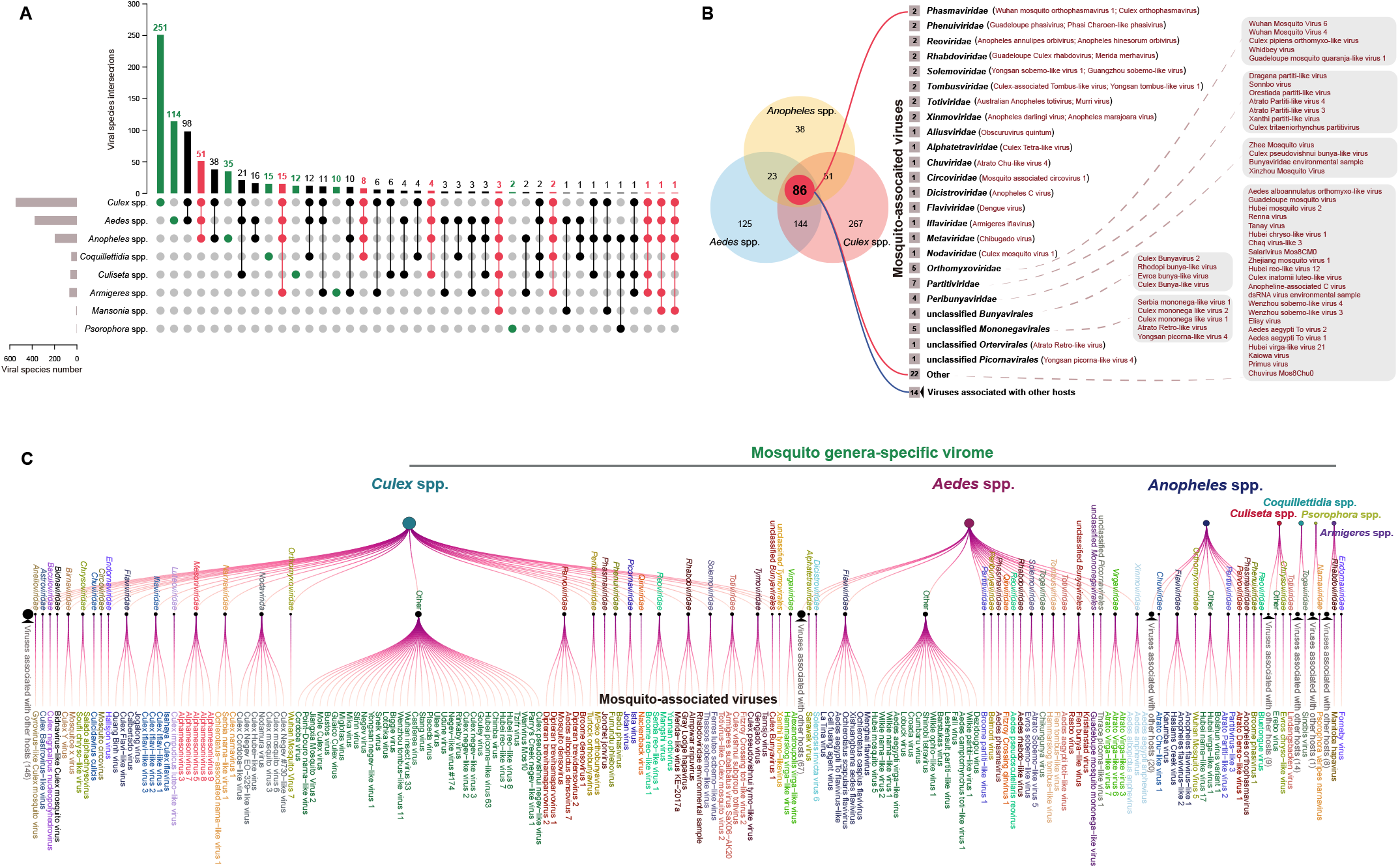
Shared and specific eukaryotic viromes in mosquitoes. (A) The upset plot showing the relationship of viromes between different mosquito genera, the number of mosquito-shared viral species among ***Aedes*** spp., ***Anopheles*** spp., ***Culex*** spp., which was marked in red. The number of mosquito-specific viral species was marked in green. (B) Shared viral species were displayed in detail through the Venn diagram. (C) Node connectivity diagram comprehensively showed the mosquito-specific viral species.

### 2.3 Core-virome in mosquitoes

Characterization and identification of core-virome of mosquitoes is imperative to understand the ecological dynamics of mosquito-associated viruses[22]. Here, we redefined the core-virome in mosquitoes at different levels of mosquito genus, continent, climate, country. Due to the heterogeneity of the number of mosquito samples at different sampling sites around the world, which could affect the frequency of eukaryotic viromes detected, we propose to analyze the core-virome with “Units”. “Units” refers to the sets of all viral species under the same province or the same county, there were a total of 83 units after such statistical adjustment. In order to find the core viruses, we summarized the distribution of the unit in *Aedes* spp., *Culex* spp. and *Anopheles* spp. continent, climate or country level. In addition, we also set the viruses that exists in more than 80% of each level as core-viome that were revealed stage-by-stage (Figure 4), To ensure the reliability, Mosquito genera and continents containing more than 10 units and climate and country containing more than 5 units were selected for further analysis.

**Figure 4.**
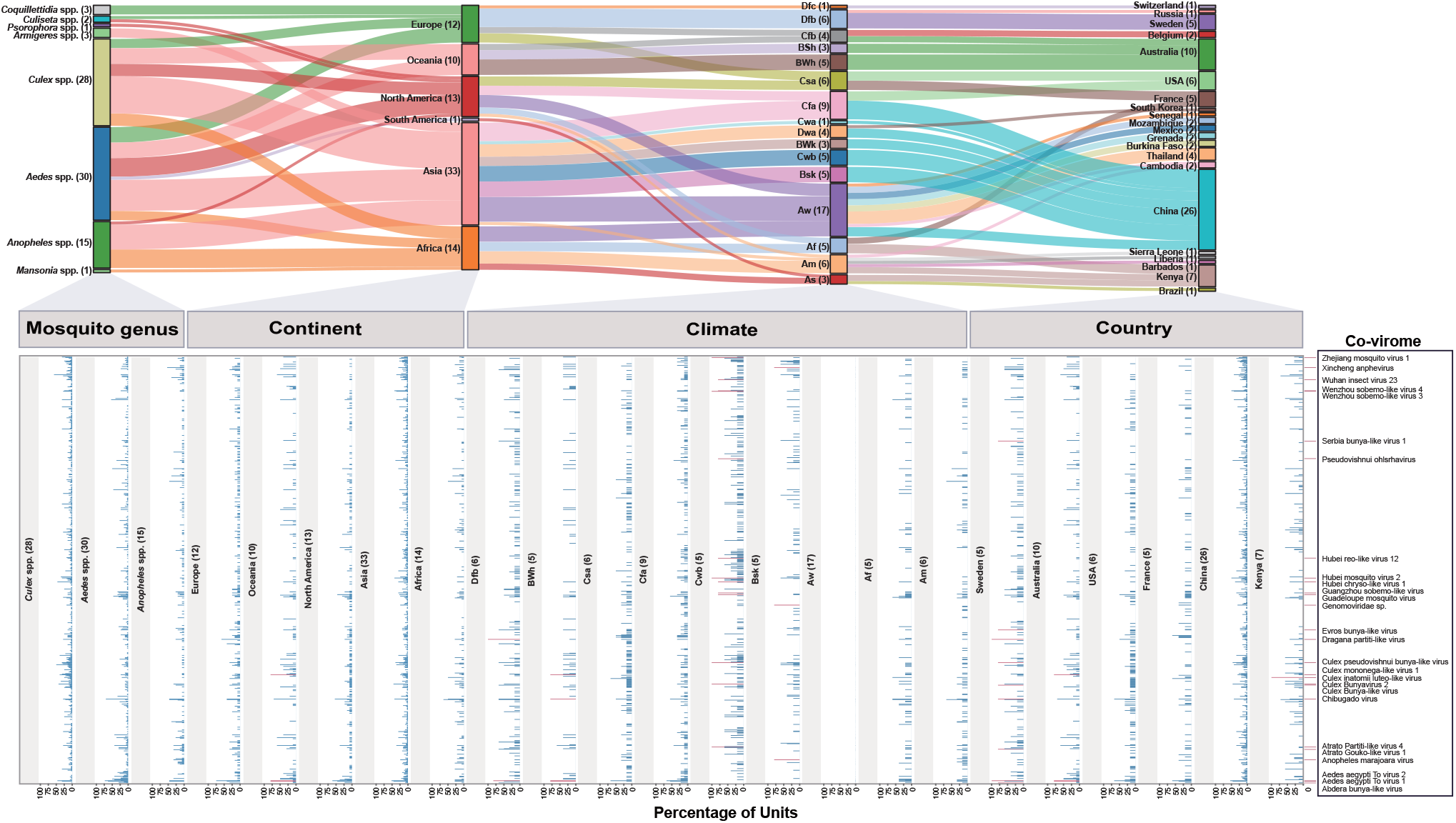
Core-virome in mosquitoes at different levels of mosquito genus, continent, climate, country is based on the ‘unit’. Sankey diagrams show the flow of units across different species, states, climates, and countries, where unit refers to the sets of all viral species under the same province or the same county; The histogram shows that the mosquito genus and continents with more than 10 units and climate and country containing more than 5 units were selected as individual element for further analysis, viruses present in more than 80% of each element are considered to be the core viruses.

It is not as expected that the three mosquito genera *Aedes* spp., *Culex* spp. and *Anopheles* spp. have no core-virome. On the continent level, Only Oceania have shared three core viruses including Aedes aegypti To virus 1, Aedes aegypti To virus 2, and Culex mononega-like virus 1. Four climate types had core viruses, the Cwb contain much higher numbers of core viruses, which were Atrato Partiti-like virus 4, Culex Bunyavirus 2, Culex pseudovishnui bunya-like virus, Hubei mosquito virus 2, Wenzhou sobemo-like virus 3, Zhejiang mosquito virus 1, Hubei chryso-like virus 1, Guadeloupe mosquito virus, Guangzhou sobemo-like virus, Hubei reo-like virus 12, Pseudovishnui ohlsrhavirus, Wenzhou sobemo-like virus 4, Wuhan insect virus 23. Dragana partiti-like virus and Aedes aegypti To virus 2 were share by Dfb, and Aedes aegypti To virus 2, Chibugado virus, Aedes aegypti To virus 1, Culex mononega-like virus 1 were share by Bwh. Anopheles marajoara virus, *Genomoviridae* sp., and Xincheng anphevirus were shared by Bsk. On the country level, Aedes aegypti To virus 2, Dragana partiti-like virus, Culex pseudovishnui bunya-like virus, Evros bunya-like virus, Chibugado virus, Culex Bunya-like virus, Serbia bunya-like virus 1, Abdera bunya-like virus, Atrato Gouko-like virus 1 were core viruses of Sweden. Aedes aegypti To virus 2, Aedes aegypti To virus 1, and Culex mononega-like virus 1 were core viruses of Australia. Only Culex inatomii luteo like virus was shared by Kenya. The results above might indicate that the co-virome was affected by environmental factors.

### 2.4 Ecological characteristic of the mosquito virome

To gain insight into the correlation of mosquito viromes with mosquito genus, climate, and location, we further investigated the key ecological driver of eukaryotic viral families by utilizing 83 Units. Pairwise comparisons of the eukaryotic viromes with ecological factors (mosquito genera, continent, climate and country) by the Mantel tests were calculated and visualized using the ggcor package at the viral families level. *Peribunyaviridae, Totiviridae*, and *Xinmoviridae* were significantly associated (0.001< Mantel’s p < 0.01) with mosquito genus. The viral families with strong correlation with collection climate is *Aliusviridae, Circoviridae, Genomoviridae*, and *Iflaviridae*. The viral families that were significantly correlated with the dual-factors of collection continent and collection country were *Phasmaviridae, Togaviridae* was strongly correlated with collection continent, and *Rhabdoviridae* and *Alphaflexiviridae* were more strongly correlated with collection country (Figure 5A). We further identified important specific viruses correlating with the mosquito genus, climate, continent, and country by the Random Forests analysis, respectively. Compared with sampling climate, continent, and country, the cross-validation of mosquito genus were smallest, indicating the limitations of mosquito genus on the distribution of viruses (Figure 5B). The representative viruses under different taxa of class were different (Supplementary Figure 7). For example, Barstukas virus, Bolahun virus variant 1 and Chibugado virus were the three most important representative viruses of mosquito genus, but the top 3 of representative viruses of collection climate were Culex Iflavi like virus 4, Genomoviridae sp. and Abdera bunya like virus. Interestingly, there were 41 shared viruses under different taxa of class (Figture 5C), which were from 16 viral families including Xincheng anphevirus, Wenzhou sobemo-like virus 4, Wuhan insect virus 23, Pseudovishnui ohlsrhavirus, Hubei mosquito virus 2, Culex pseudovishnui bunya like virus, Culex Bunyavirus 2, Culex mononega like virus 1, Anopheles marajoara virus and Abdera bunya like virus (Figture 5D). These results indicated that the distribution of these shared viruses may be affected by the combination of mosquito species, sampling sites and climate.

**Figure 5.**
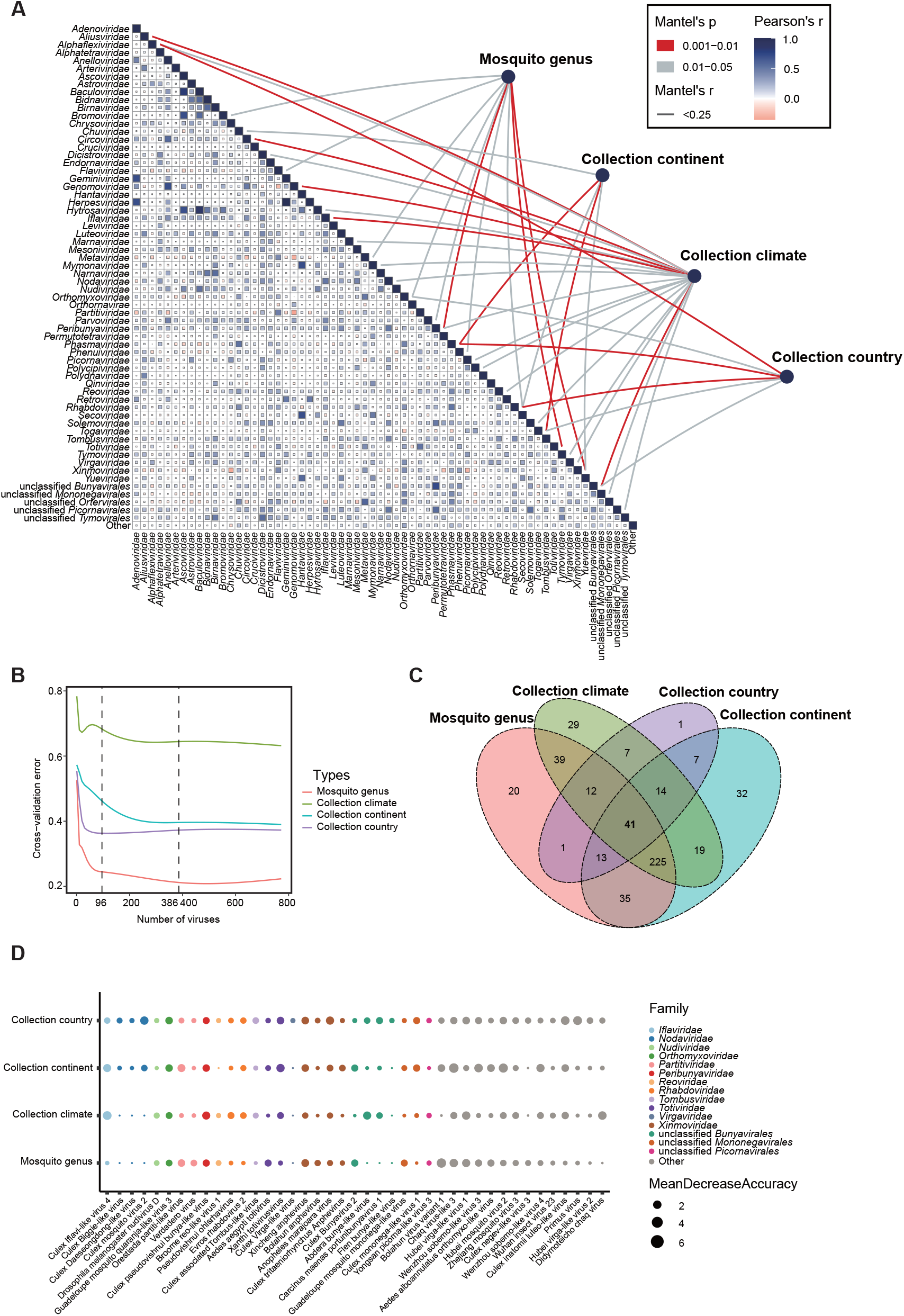
Ecological drivers of the eukaryotic viromes composition in the mosquitoes. (A) Pairwise comparisons of the eukaryotic viromes with ecological factors (mosquito genera, continent, climate and country) by the Mantel tests, which was shown with a color gradient denoting Spearman’s correlation coefficient. Edge width corresponds to the Mantel’s r statistic for the corresponding distance correlations, and edge color denotes the statistical significance based on 9,999 permutations. (B) The line chart displaying the 10-fold cross-validation error changes with the number of viruses under the mosquito genus, climate, continent, and country, which was used to identify the marker taxa according to the parsimony principle. The dotted line indicating the number of viruses chosen when the cross-validation error was stabilized; (C) Venn diagram showing intersection of the predicted important viruses among the mosquito genus, climate, continent, and country; (D) Dotplot shows the most representative tagged viruses at the level of mosquito genus, climate, continent and country.

### 2.5 The marker viruses reveal the migration events of mosquitoes

Although mosquitoes can fly only a limited distance, passing natural winds or attaching themselves to vehicles may help them spread further. To explore the cross-regional migration of mosquitoes, we sought to select some viruses from the identified core-virome that are prevalent in several regions as marker viruses to characterize mosquito movement. By calculating the relative abundance of each virus, we found that Aedes aegypti To virus 2, Chibugado virus, Phasi Charoen-like phasivirus, and Wenzhou sobemo-like virus 4 were highly expressed (top 20) in at least 3-4 continents and had potential as marker viruses (Supplementary Figure 8). We then further analysed the ancestral geographical regions and migration patterns of these viruses. The results showed that the Aedes aegypti To virus 2 carried in mosquitoes from China, Senegal, Liberia, Brazil, and USA originated from Australia, in addition to the close evolutionary relationship between the Chinese strain and isolates from Cambodia, Sweden, Belgium, Senegal, and Brazil, suggesting that mosquitoes are reciprocally transmitted between these regions (Figure 6A). Chibugado virus originated in Senegal and spread to Australia, China and Cambodia, with the Australian strain also clustered with isolates from USA, Colombia and Switzerland. The other two marker viruses originated in Asia, with Phasi Charoen-like phasivirus spreading from Thailand to India, Australia and Brazil, and the Australian isolate had a close evolutionary relationship with the Kenyan isolate; Wenzhou sobemo-like virus 4 originated in China and spread to Spain, Switzerland and the USA (Figure 6A).

**Figure 6.**
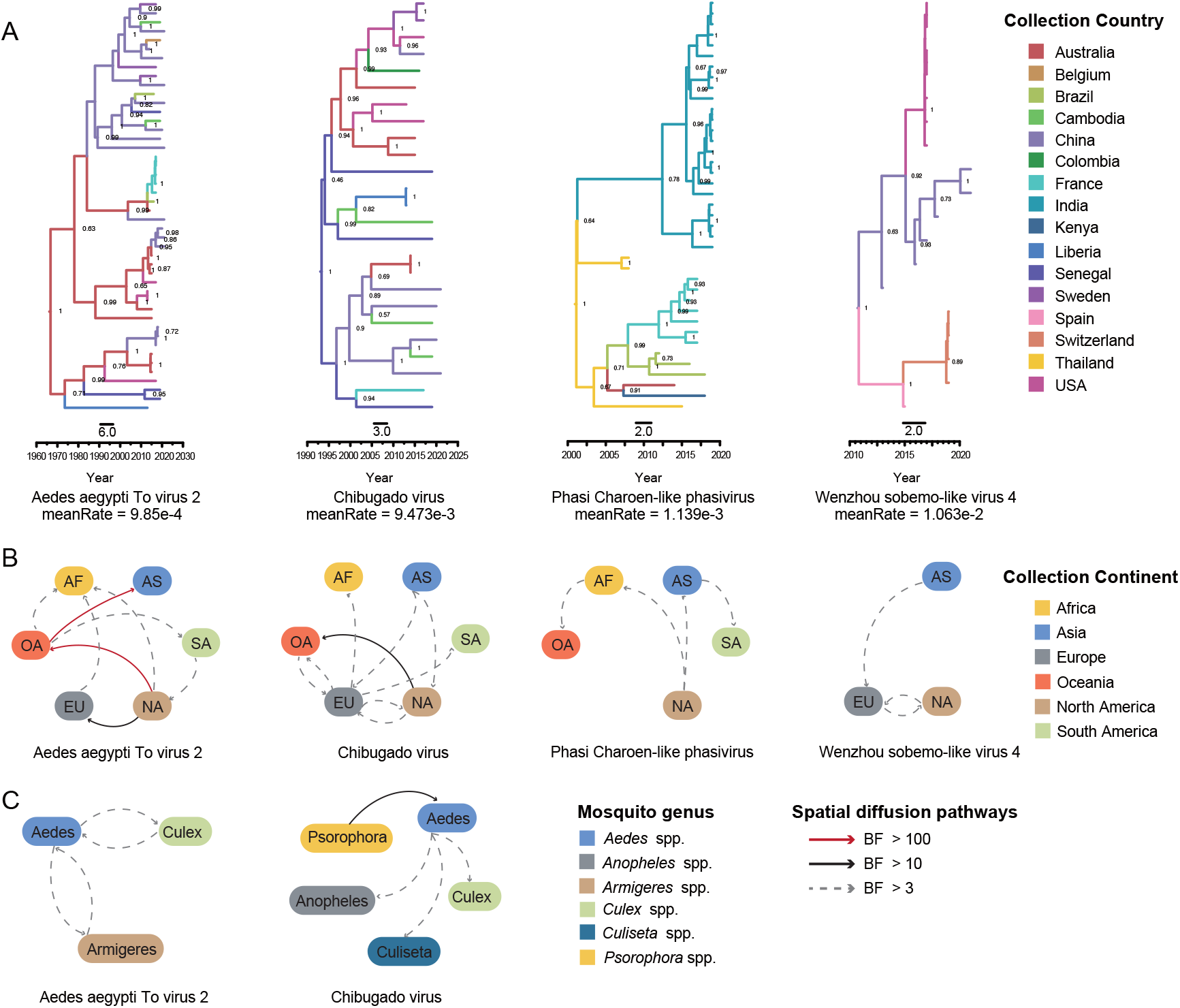
Spatial diffusion pathways of potential marker viruses in mosquitoes. Mosquito-borne viruses that are prevalent across multiple areas were used as markers to track the movement of mosquitoes. The geographic transmission routes of Aedes aegypti To virus 2 (n = 46), Chibugado virus (n = 25), Phasi Charoen-like phasivirus (n = 39), and Wenzhou sobemo-like virus 4 (n = 18) were reconstructed using BEAST v1.10.4. The datasets include all available sequences in GenBank with sampling information, and contigs assembled from NGS data. (A) MCC tree based on partial region of RdRp gene of potential marker viruses. The color of the branches represents the sampling country, and only the posterior of nodes larger than 0.6 are shown. B-D, Diffusion pathways of marker viruses across continents (B) and mosquito genera (C) with BF > 3 are shown.

The transmission of four candidate marker viruses across continents, mosquito genera of host, and climates was assessed by BSSVS analysis. The results showed a total of 17 diffusion pathways of marker viruses in transcontinental transmission, of which three pathways were strongly supported, including Oceania to Asia (BF=780), North America to Oceania (BF=400), and North America to Europe (BF=39) (Figure 6B). There were varying degrees of overlap in the diffusion pathways of the four marker viruses, with six pathways being significantly supported in more than two marker viruses, including Asia to Europe, Europe to Africa, Europe to North America, North America to Europe, North America to Africa, and North America to Oceania. We found more frequent mutual transmission of viruses between North America, Europe, Oceania and other regions, suggesting that they may play a key role in the transcontinental migration of mosquitoes. In addition, there was no significant evidence to support the communication of mosquitoes between Africa and South America and Asia. To further explore which mosquito genera play a vector role in the transmission of marker viruses, host information on isolates was collected and the BSSVS procedure was implemented. The results showed strong support for transmission from the *Psorophora* spp. to the *Aedes spp*. (BF=12). Moreover, the *Aedes* spp. is associated with a variety of mosquito genera for viral transmission (Figure 6C). Finally, considering that climates is also a non-negligible factor influencing mosquito activity, we likewise analysed the spread of marker viruses between different climatic zones. A total of 39 mosquito migration routes were identified, but only one route, Csa to Cwa, was supported in more than one marker viruses (Supplementary Figure 9).

## 3. Discussion

Viral metagenomic studies have driven the explosion of mosquito-associated viruses discovery, which not only contributes to detection of arboviruses, more to overview of viruses circulating in mosquitoes[23]. These studies have shown the diversity and abundance of virome in *Culex* spp., *Aedes* spp., *Mansonia* spp., *Armigere* spp., *Psorophora* spp., *Coquillettidia* spp., *Culiseta* spp., and *Anopheles* spp. [24–29], but localized in specific regions. Nevertheless, there is still a lack of broader studies that directly compare mosquito virome from large multisite. In our study, we integrated global metagenomic datasets of mosquitoes, and demonstrated “pan-virome” that we defined as total viruses in overall mosquito, the “pan-virome” include 773 eukaryotic viral species. Further, we analyzed the distribution of features at different mosquito genus for “pan-virome”. Among 8 mosquito genus, *Culex* spp., *Aedes* spp., and *Anopheles* spp. had enriched “shared-virome”, which illustrates the great potential of these viruses for viral circulate between different mosquito genera. 7 mosquito genus were observed the genera-specific virome in addition to the *Mansonia* spp. the mosquito-genera specific virome could have been behavior-related like the differences of hematophagous and feeding habits in adult mosquitoes, or the influence of environmental factors, especially the preference of the aquatic environment in which the larvae live. Moreover, we profiled the dominant viral families among different mosquito genus. *Phasmaviridae* is the dominant viral family with high abundance in *Culex* spp., *Aedes* spp, and *Anopheles* spp., and the *Coquillettidia* spp. contains the most prevalent *Picornaviridae* and the Unclassified *Picornavirales*. It is still noteworthy that the *Flaviviridae* was highly abundant in *Anopheles* spp.

The concept of core-virome emerged with the propose of commensal microbiome that, any set of viral taxa in most cases, as well as the associated functional attributes characteristic of a specific host or environment[30]. Comparison of core-virome among mosquito host/ environment in a habitat is central in research of ecological dynamics of mosquito-associated viruses[22]. Core-virome are typically quantified by the occurrence of viruses across multiple samples[16,17,22], but the choice of spatial scale as well as sampling number could have a strong influence on the composition of the core[30]. Here, we present the “Unit” that refer to the occurrence of viruses across the province or county. There are several previous studies have shown the core-virome seems to characterize the individual mosquito species, and whether the virome of a species differs significantly between different ecological niches have not resolved. We performed analytical analyses of core-virome from the perspective of mosquito genus, continent, climate and country, more core-virome were found associated with environmental factors, which may signify transmission and adaptation of viruses between ecological niches.

The host and environment are critical significant selective territory that not only contributes to the adaptation of viruses, but also affects the diversity and distribution of viruses. Thus, integrating viral metagenomics into an ecological framework is very necessary to address rarely considered ecological questions in virology, such as the limitations of viral host and geography, which may increase understanding of viruses from other dimensions and explain how they interact with their surroundings. Japanese encephalitis virus (JEV), a member of the genus *Flavivirus*, is endemic to Southeast Asia and China and humans are mainly infected via the bite of infected *Culex spp*, and as a natural reservoirs and amplifying host for it[31,32]. The vectors of JEV depends on the abundance of host in each location in India, suggesting the host selection depending on the densities of available vertebrate hosts in a given location [33]. In addition, the transmission of arboviruses may be affected due to ecogeographic barriers, currently little has been revealed. Mosquitoes, as one of the most important vectors of arboviruses, play an critical role in the transmission of these viruses. Here, we comprehensively analyzed the correlation between the eukaryotic viromes with ecological factors (mosquito genera, continent, climate and country) using 83 Units and found that some viral families were greatly correlated with specially designated ecological factor (Figure 5). No virus is related to all these ecological factors at the same time. Also, the top 30 of specific representative viruses under different taxa were not the same (Supplementary Figure 7). We identified the 41 shared important viruses under different taxa of class using Random Forests, and these viruses have 5 overlap viruses with the the previously obtained core-virome, including Anopheles marajoara virus, Culex Bunyavirus 2, Hubei mosquito virus 2, Pseudovishnui ohlsrhavirus, Wuhan insect virus 23, and Xincheng anphevirus, which may play a very important role in viral ecology.

Both mosquito and bird are carriers of numerous viruses, the movement routes of mosquitoes are not as clear studied as those of wild birds, which is determined by their ability to fly - only within a limited habitat. However, in the presence of external interventions, mosquitoes’ cross-regional movements may bring original area-restricted viruses to new areas and cause outbreaks, such as the 2014 dengue fever outbreak in Guangdong caused by imported cases. On the other hand, in the context of global climate change, mosquitoes that prefer warm and humid climates may migrate in search of new, more livable environments. All these possible mosquito activities will pose a risk to the cross-regional transmission of mosquito-borne viral infections. In this study, we tried a new strategy to track the activities of host mosquitoes by analyzing the transmission routes of several viruses commonly carried by mosquitoes as markers. Considering the viral relative abundance of each sample, we did not use the previous five viruses at the Units level above, but chose another four viruses: Aedes aegypti To virus 2, Chibugado virus, Phasi Charoen-like phasivirus, and Wenzhou sobemo-like virus 4 as potential marker viruses to tracing the mosquito migration, three of which were included in the previously obtained core-virome. The results showed that mosquitoes in North America, Europe, and Oceania have more frequent cross-regional communications (Figure 6). These continents have more developed and mature tourism and transportation resources, which may be one of the reasons for the cross-regional activity of mosquitoes. It should be noted that due to the wide variety of climatic zones and limited available sample data, it is difficult to obtain uniform results across marker viruses (Supplementary Figure 9), thus cannot accurately elucidate the transmission characteristics of mosquitoes among different climates. In the future, researchers from more regions may need to work together to compensate for the regional balance of samples by using a standardized sampling process and appropriate sequencing depth, so as to obtain more reliable evidence of mosquito migration.

In conclusion, we systematically and clearly describe a global map of mosquito virome diversity, spatial structure, and its ecological determinants based on the existing mosquito metavirome data including public and our laboratory acquired data, which will provide a more comprehensive view of the mosquito virome than previous studies, and inspire us to look at the virus composition, interaction and evolution of mosquito hosts from the perspective of the whole virus, which also may provide help for virus prevention and monitoring in the future.

## 4 Materials and Methods

### 4.1 Mosquito collection

The mosquito samples were collected from different provinces including Guangdong, Xinjiang, and Hubei in China during the period 2014-2021. After species identification based on morphological features using morphology descriptions, the species of mosquitoes belonged to three genera and eight species containing *Aedes albopictus, Aedes vexans, Aedes flavescens, Aedes caspius, Culex pipien, Culex quinquefasciatus, Anopheles Hyrcanus* and *Anopheles sinensis*. In addition to these field mosquito samples, the samples from different development stages (egg, larvae, pupae, and adult) bred in the lab in 2017 were also collected, which was originated from the Chinese Center for Disease Control and Prevention. The mosquitoes were stored at -80 ° or on dry ice until processed.

### 4.2 RNA extraction and sequencing

The mosquitoes were pooled (about 50 individuals per pool) by the species, gender, collection site and date. The mosquitoes of one pool were mixed in one tube and the electric tissue grinder (Tgrinder QSE-Y30) was used to grind with a pestle (without RNase) on ice after 200 μL of RPMI medium was added to each tube. The homogenate of the mosquitoes was centrifuged at 20,000 x g for 30 min at 4 ° and then filtered by a 0.45 μm pore size filter with most bacterial or tissue debris removed. Subsequently, the supernatant was collected and used for the extraction of viral RNA using the QIAamp Viral RNA Mini Kit according to the operation manual (https://www.qiagen.com/cn/shop/automated-solutions/qiaamp-viral-rna-mini-kit/). After the viral RNA was extracted, the TruSeq® Stranded Total RNA Sample Preparation Kit (Illumina, America) was used to construct the sequencing libraries. Qualified libraries were pooled to the flowcell according to the requirements of effective concentration and target data volume, and paired-end sequencing was performed using Illumina HiSeq 2500 sequencing platform.

### 4.3 Public data collection

The viral metagenomic data of mosquitoes was downloaded from Gene Expression Omnibus (GEO, https://www.ncbi.nlm.nih.gov/geo/). Meanwhile, the data were also searched from the published articles related to mosquito viral metagenomics. To ensure the availability and accuracy of follow-up analysis, the data of mixed mosquitoes were removed. Finally, the 608 data were selected for analysis (see Supplemental Table 1). The sample information was obtained from the SRA Run Selector (https://www-ncbi-nlm-nih-gov.ezproxy.u-pec.fr/Traces/study/?) or the related published articles. The climate of the sampling site was queried from the Weather and Climate website (https://tcktcktck.org).

### 4.4 Viral metagenomic analysis

After the raw data obtained by RNA sequencing was quality controlled using Trim Galore v0.6.6 with the adapter and low-quality reads removed, the clean reads were de-novo assembled by Trinity v2.1.1 and Quast v5.0.2 was subsequently used to assess the quality of the assembled contigs. The longest transcripts were reserved after redundancy was removed using the built-in de-duplication scripts in Trinity according to the naming rules. The assembled contigs were annotated using a BlastX search against the NCBI non-redundant (nr) sequence database on sensitive mode using Diamond with the best hits and an e-value cutoff of < 10-5. To further verify the results of this annotation, the clean reads were also classified by Kraken2 with the standard Kraken2 database containing the refseq bacteria, archaea, and virus, whose results were interpreted by Pavian (https://fbreitwieser.shinyapps.io/pavian/) website. The contigs mapped to the virus were extracted with the length > 500bp. The virus abundance was assessed by mapping the clean reads to these assembled viral contigs using the BBmap software. The ComplexHeatmap package was applied to view the distribution of viruses in different samples. The intersections and unions of viruses between different genera of mosquitoes were shown by the UpSetR package and VennDiagram package. The distance correlations between genomic and ecological environmental data were performed using the ggcor package. Next, the relative abundance of all viruses carried in the samples was calculated and visualized using R v4.1.1, the results of the top 20 viruses with the highest abundance were retained and further classified by sampling continent to calculate the mean abundance. Viruses expressed in high abundance in at least 3 continents were further used as candidate marker viruses to track the migration routes of mosquitoes.

### 4.5 Screening of eukaryotic virus

The host range of mapped viruses was searched from NCBI based on the species names. The viruses related to fungi, bacteria, and protozoa were filtered out according to the host of viruses. The host of remaining viruses was classified, which was visualized by the ggraph package.

### 4.6 Important viruses predicted using Random Forests

The important viruses were identified at the unit level under the mosquito genus, climate, continent, and country by the random forest algorithm using the randomForest package with default parameters. In addition, the number of the important viruses were obtained using the 10-fold cross-validation implemented in the rfcv function of the randomForest package with five repeats based on the parsimony principle.

### 4.7 Geographical reconstruction of marker viruses transmission

All RdRp gene sequences of four marker viruses available in GenBank were collected and merged with the assembled contigs for multiple sequence alignment, and the multiple sequence alignment files were further trimmed according to the reference sequences (MT913596.Aedes aegypti To virus 2 isolate AaTV2/BR\_TO: 2461-3006, MN661043.Chibugado virus strain Psal 1744-4: 4403-4948, NC\_038262.Phasi Charoen-like virus isolate Rio segment L: 3208-4059, NC\_033138.Wenzhou sobemo-like virus 4 strain mosZJ35391: 1699-1974), respectively. After removing sequences of samples with unknown sampling year and location, four smaller datasets, Aedes aegypti To virus 2 (n = 46), Chibugado virus (n = 25), Phasi Charoen-like phasivirus (n = 39), and Wenzhou sobemo-like virus 4 (n = 18) were subjected to discrete trait ancestral reconstruction using BEAST v1.10.4[34]. After determining the collection time and location corresponding to the sequences, the HKY substitution model under the uncorrelated relaxed clock was selected with 500 million iterations, and Coalescent Constant Population was used as the prior model. Tracer v1.7.2 was used to read and detect the results of numerical simulations, and the results were considered credible for the next analysis only when the ESS values of all parameters are greater than 200. The tree files were summarized by TreeAnnotator with 10% burn-in cutoffs and visualized by FigTree (http://tree.bio.ed.ac.uk/software/figtree/). The climate type, and mosquito genus of the samples were analyzed as separate traits for Bayesian stochastic search variable selection (BSSVS). Whether the propagation was significant was determined based on Bayes factor (BF), where 100 ≤ BF < 1,000 indicates very strong support, 10 ≤ BF < 100 indicates strong support, and 3 ≤ BF < 10 indicates statistically significant support. The spatial diffusion pathways were visualized by SpreaD3 v0.9.6 [35] and Adobe Illustrator CC 2022.

### 4.8 Data and code availability

The raw sequencing data from the lab reported in this paper have been deposited in the Genome Sequence Archive (GSA) of the National Genomics Data Center (https://ngdc.cncb.ac.cn), Beijing Institute of Genomics (BIG), Chinese Academy of Sciences, under accession numbers HRAxxxxxx that can be accessed at https://bigd.big.ac.cn/gsa/HRAxxxxxx. All other original data were available from the corresponding authors on request.

